# Discovery and characterization of small molecule inhibitors of CBL-B that act as intramolecular glue to enhance T-cell anti-tumor activity

**DOI:** 10.64898/2026.01.22.701125

**Authors:** Gajewski Stefan, Taherbhoy M. Asad, Boyle Kathleen, Gosling Jennifa, Perkins Jilliane, Dhamnaskar Ketki, Sheung Julie, Baker Kenton, O’Connell Nichole, Bravo Brandon, Mukerji Ratul, Tenn-McClellan Austin, Kurylo Katherine, Haria Dhwani, Gallotta Marilena, Juan Joseph, Tan May, Prakash Sumit, Weiss R. Dahlia, Lawrenz Morgan, Cardozo Mario, Wang Chenbo, Cummins Thomas, Clifton C. Matthew, Rountree Ryan, Saha Anjanabha, Zapf W. Christoph, Hansen M. Gwenn, Sands T. Arthur, Cohen Frederick

**Affiliations:** Nurix Therapeutics

**Keywords:** Immuno-oncology, small molecule drug discovery, molecular glue, T-cell activation

## Abstract

CBL-B is a RING-type E3 ubiquitin ligase that acts as a critical negative regulator of T-cell activation. It promotes T-cell anergy and suppresses immune responses through ubiquitin-mediated control of signaling proteins at the immunological synapse. T cells deficient in CBL-B activity lose their dependence on CD28 co-stimulation, exhibit heightened activation and increased cytokine production, and fail to re-establish anergy. In addition, mice deficient in CBL-B activity reject tumors. Together, this cellular mechanism and *in vivo* phenotype suggest inhibition of CBL-B may be a viable immuno-oncology therapeutic strategy. Here, we report the rational design and execution of a high-throughput screen (HTS) to identify small molecule inhibitors of CBL-B. This campaign led to the discovery of a scaffold that inhibits CBL-B E3 ligase activity with micromolar potency. Structural characterization revealed an intramolecular glue mechanism, in which the compound stabilizes the closed state of CBL-B, preventing phosphorylation of a tyrosine residue that is critical for activation and E2 binding. Iterative structure–activity optimization yielded compounds with nanomolar activity that enhanced T-cell activation and cytokine secretion in primary human T cells and suppressed tumor growth in a syngeneic colorectal mouse model. Together, these studies validate the biological rationale for pharmacological CBL-B inhibition and enabled the *de novo* discovery of intramolecular CBL-B glue inhibitors. This work culminated in the identification of NX-1607, a first-in-class oral CBL-B inhibitor now in clinical development for cancer immunotherapy.

## Introduction

Peripheral T-cell activation is tightly regulated to enable adaptive responses to immunogens while preventing potential harm (cytotoxicity or autoimmunity) to the host’s own tissues. Casitas B-lineage lymphoma proto-oncogene B (CBL-B), a RING finger E3 ubiquitin ligase of the CBL family, serves as a key regulator of immune activation and peripheral tolerance (1). CBL-B is predominantly expressed in peripheral T cells and associates with cytosolic components of the T-cell receptor (TCR) and other membrane receptor signaling cascades, where it downregulates their activity by ubiquitylating downstream proteins (2, 3). Loss of CBL-B uncouples the requirement for CD28 co-stimulation to enable T cell proliferation and IL-2 production indicating that CBL-B is involved in the CD28 costimulatory signaling pathway (4, 5). CBL-B KO mice show spontaneous tumor rejection and improved survival in preclinical tumor models (6). Similarly, adoptively transferred CBL-B-deficient CD8+ T cells show enhanced tumor infiltration/persistence, resist immunosuppression (Tregs, TGFβ or PDL-1), secrete more IFN-γ, and exhibit significantly improved anti-tumor efficacy (5, 7–9). A 2011 study demonstrated that mice with a point mutation in the CBL-B RING domain that disrupts E2 binding have a T cell phenotype similar to knockout animals and spontaneously reject tumor cells (8). Taken together, these data support the concept that pharmacologic inhibition of CBL-B ligase activity can lower the activation threshold of T cells and drive stronger antitumor immune responses, providing a rationale for pursuing CBL-B as a drug target in immuno-oncology.

The role of CBL-B in T-cell adaptive immunity is primarily to restrain TCR signaling by applying specific ubiquitin modifications to key signaling proteins through both proteasome-dependent and nonproteolytic mechanisms (1, 2, 9–11). K33-linked polyubiquitylation of the TCR-ζ chain at residue K54 reduces TCR-ζ phosphorylation and decreases both the association with, and phosphorylation of, ZAP70 (12, 13). CBL-B polyubiquitylation of the p85 subunit of PI3K disrupts its association with CD28 and TCR-ζ, suppressing co-stimulatory activity (5, 14–17). Additional key targets of CBL-B include LAT, a scaffolding protein critical for T-cell activation (18), the guanine nucleotide exchange factor VAV1 (19), and the phospholipase PLCG1 (20), which plays a key role in calcium release downstream of TCR signaling. The levels of CBL-B protein itself are regulated through CD28 signaling via PKCθ-mediated phosphorylation and subsequent NEDD4 dependent proteasome directed polyubiquitylation (11, 21).

Expression of CBL-B is induced by sustained TCR/CD3 signaling and CTLA-4 costimulatory signaling (22) while PD-L1 silencing reduces CBL-B expression in CD8^+^ T-cells (23). These findings identify CBL-B as a pivotal intracellular regulator that fine-tunes the costimulatory threshold governing T-cell activation. By modulating the balance between stimulatory inputs, CBL-B establishes a defined activation window in which the lower boundary is dictated by TCR/CD3 engagement, while the upper boundary is constrained by CD28-mediated signaling. Notably, under conditions of CD3-only stimulation, loss of CBL-B results in an activation level comparable to that observed during CD3/CD28 co-stimulation of CBL-B competent T cells, whereas CD3/CD28 co-stimulation in the absence of CBL-B leads to pronounced hyperactivation (8).

The domain architecture of CBL-B comprises an N-terminal tyrosine kinase binding (TKB) domain, containing a four-helix bundle (4H), an EF-hand (EFH), and an SH2 domain that form a globular fold (24). The TKB domain contains the degron recognition motif and binds to degrons containing a phosphotyrosine residue. A short linker helix region (LHR) separates the SH2 from the RING domain, which recruits ubiquitin-charged E2 enzymes. In the closed, inactive state, Y363 within the LHR is positioned between the SH2 and EFH domains, thus tethering the RING domain to the TKB fold. Downstream of the RING domain (residues 427-982), the protein is a predominantly unstructured C-terminal region that contains phosphorylation sites, proline-rich sequences, and a ubiquitin-associated (UBA) domain that mediates interactions with SH2- and SH3-domain-containing partners and ubiquitylated proteins. This modular architecture enables it to function as both a scaffold in tyrosine phosphorylation-regulated receptor complexes and a ubiquitin ligase (2, 25).

CBL-B activation begins with binding of a phosphotyrosine-containing peptide to the SH2 domain, displacing a F263 containing loop into the LHR-binding groove (Figure 3A) (24). This dislodges the LHR helix from the TKB, yielding a transient open, inactive state that resolves via either LHR rebinding or Y363 phosphorylation. Phosphorylation stabilizes the open, active state, freeing the RING domain to engage E2 enzymes and ubiquitylate substrates (26). In the open state, the LHR acts as a flexible linker, and Y363 phosphorylation blocks TKB rebinding but is reversible by SHP-1 dephosphorylation (27).

This activation mechanism suggested two potential inhibition strategies: stabilizing the closed state with unphosphorylated Y363 sequestered in the TKB or blocking E2 interactions in the open Y363 phosphorylated state. We developed time-resolved fluorescence resonance energy transfer assays (TR-FRET) for the closed and open states of CBL-B to support high throughput screening (HTS) campaigns to identify small molecule inhibitors. Although both modalities were screened, only screening against the closed state yielded viable hits. Initial hits were validated using dose-response FRET assays and label-free surface plasmon resonance (SPR). Following triage, a single compound showed reproducible, dose-dependent CBL-B binding across assays. Co-crystallization of this molecule with CBL-B revealed its absolute configuration, the binding pose, and its inhibitory mode as an intramolecular glue stabilizing the closed state. Leveraging validated assays, we launched a structure-based design effort that yielded potent analogs suitable for studying CBL-B inhibition in cells. These inhibitors induced dose-dependent upregulation of T-cell activation markers and cytokine secretion in human peripheral blood mononuclear cells (PBMCs). When administered orally to mice bearing syngeneic tumors, an early analog of the CBL-B inhibitor series reproduced the knockout phenotype (8), resulting in enhanced IL-2 secretion, increased surface marker expression, and tumor growth inhibition. Further optimization for potency and pharmacokinetics resulted in the discovery of NX-1607, a potent CBL-B inhibitor currently being tested in clinical trials (28).

## Results

### High-Throughput Screen (HTS) Campaign to Identify CBL-B Inhibitors

To identify CBL-B inhibitors, we developed a homogeneous time-resolved fluorescence (HTRF)-based HTS assay to monitor the transition between the closed inactive and open activated conformations. In this assay, CBL-B phosphorylation was catalyzed by Src kinase in the presence of a phosphotyrosine-containing peptide derived from ZAP70, which acts as a coactivator to promote the open conformation. Phosphorylation at Y363 was detected via fluorescence resonance energy transfer (FRET) between a terbium-labeled anti-phosphotyrosine antibody and a streptavidin-conjugated XL665 fluorophore bound to the biotinylated N-terminus of CBL-B (Figure 1 A).

**Figure 1:**
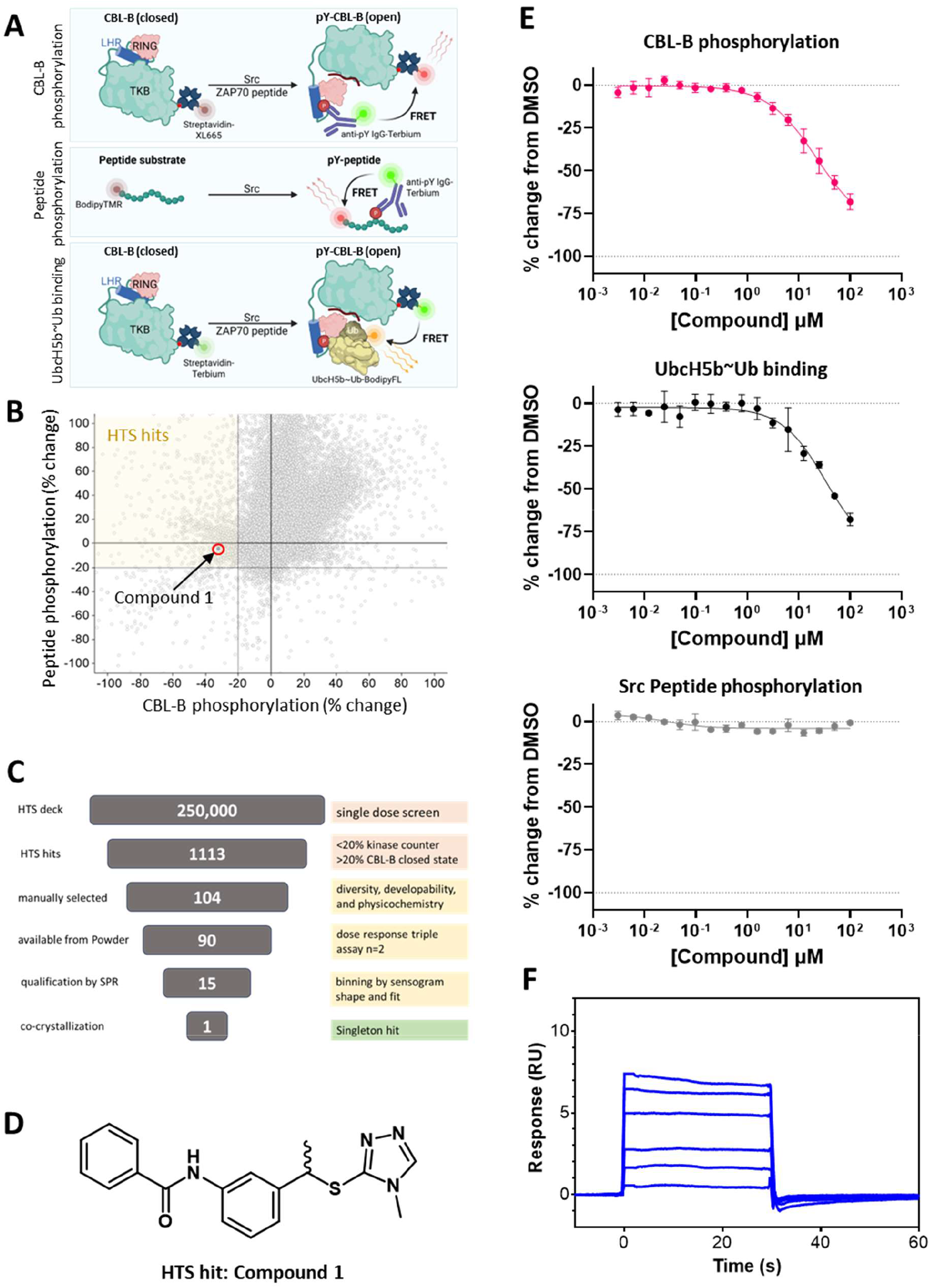
High throughput screen to discover small molecule inhibitors of CBL-B. A) Schematic of the HTS assays designed to detect Src-mediated phosphorylation of CBL-B or an RTK peptide. Phosphorylation of CBL-B was detected using a phosphotyrosine-specific antibody targeting Y363, which generated a FRET signal with a fluorophore attached to the N-terminus of CBL-B. In the peptide counter-screen assay, phosphorylation of the RTK peptide was similarly detected using the same antibody, producing a FRET signal with a fluorescent dye at the peptide’s N-terminus. Compounds that inhibited CBL-B phosphorylation without affecting Src activity in the peptide assay were presumed to stabilize CBL-B in its closed conformation. An additional FRET assay monitoring UbcH5B~Ub binding to phosphorylated CBL-B was used to confirm that the HTS hits bind to the inactive, closed state of CBL-B. B) Scatter plot of the phosphorylation HTS performed against ~250,000 compounds. Each point represents a single compound, plotted by percent change of CBL-B phosphorylation (x-axis) and percent change of RTK peptide phosphorylation (y-axis). Solid lines at y = 0 and x = 0. indicate no inhibition. Dotted lines at x = –20 and y = –20 mark percent inhibition cutoffs. The shaded area x < −20 and y > −20 percent change indicates the region of active HTS hits. C) Hit validation funnel. Numbers in the central boxes denote the number of compounds remaining at each stage of the cascade (primary HTS, retest, counter screen, dose–response, biophysical validation), culminating in a single confirmed hit. D) Chemical structure of the validated HTS hit, compound **1**, identified as a racemic mixture. E) Dose response curves for compound **1** in the CBL-B phosphorylation assay, the UbcH5B~Ub binding assay, and the Src peptide phosphorylation counter screen. F) SPR sensograms for compound **1**, tested at a top concentration of 50 μM with a two-fold serial dilution. Equilibrium analysis yielded a K_D_ of 48 µM.

To ensure specificity and exclude false positives arising from Src inhibition, we implemented a counter screen using a generic receptor tyrosine kinase (RTK) substrate peptide phosphorylated by Src kinase. In this assay, FRET signal was generated between the same terbium-labeled antibody and a BODIPY-TMR fluorophore conjugated to the peptide’s N-terminus (Figure 1 A).

A library of over 250,000 small molecules was screened at a single concentration of 25 µM. Primary hits were defined as compounds that produced a ≥20% reduction in FRET signal in the CBL-B phosphorylation assay compared to vehicle-treated controls, while exhibiting ≤20% reduction in the Src counter screen assay (Figure 1B). Hits meeting these criteria were clustered and analyzed using cheminformatic tools to assess chemical diversity, physicochemical properties, and overall chemical developability. Based on this analysis, a representative set of 104 compounds was selected, and 90 compounds with available fresh powder stocks were obtained to generate full dose–response curves (Figure 1 C).

An additional FRET assay was developed to directly measure binding of ubiquitin-charged E2 ubiquitin-conjugating enzyme (UbcH5B~Ub) to the active, open state of CBL-B, adding a further layer to ensure that hits specifically inhibit CBL-B activation. Compounds that block activation are expected to stabilize the autoinhibited closed state of CBL-B, thereby preventing E2 recruitment. In this assay, CBL-B was labeled with a streptavidin–terbium donor fluorophore, and binding of BODIPY-FL–labeled UbcH5B~Ub to the RING domain was monitored via FRET (Figure 1 A). Activation of CBL-B was achieved as in the CBL-B phosphorylation assay through addition of a ZAP70-derived phosphotyrosine-containing peptide and Src kinase. Full dose-response curves of the 90 hits were generated using all three assays in parallel.

Fifteen compounds that exhibited inhibition of both CBL-B phosphorylation and E2 binding, without detectable inhibition of Src kinase activity, were advanced for further evaluation. These compounds were assessed by SPR to measure direct binding to CBL-B. The screening cascade led to the identification of compound **1** (Figure 1 D), which inhibited CBL-B activation with IC_50_ values of 26 μM and 29 μM in the phosphorylation and E2 binding HTRF assays, respectively. No Inhibition of Src kinase activity was observed (Figure 1 E, Table 1). Importantly, compound **1** was the only hit to show orthogonal binding to CBL-B in the SPR assay, with an apparent dissociation constant (K_D_) of 48 μM (Figure 1 F, Table 1), qualifying it for further structural characterization and optimization. The data confirm specific inhibition of the closed state of CBL-B with modest potency and minimal effect on Src kinase activity.

**Table 1:**
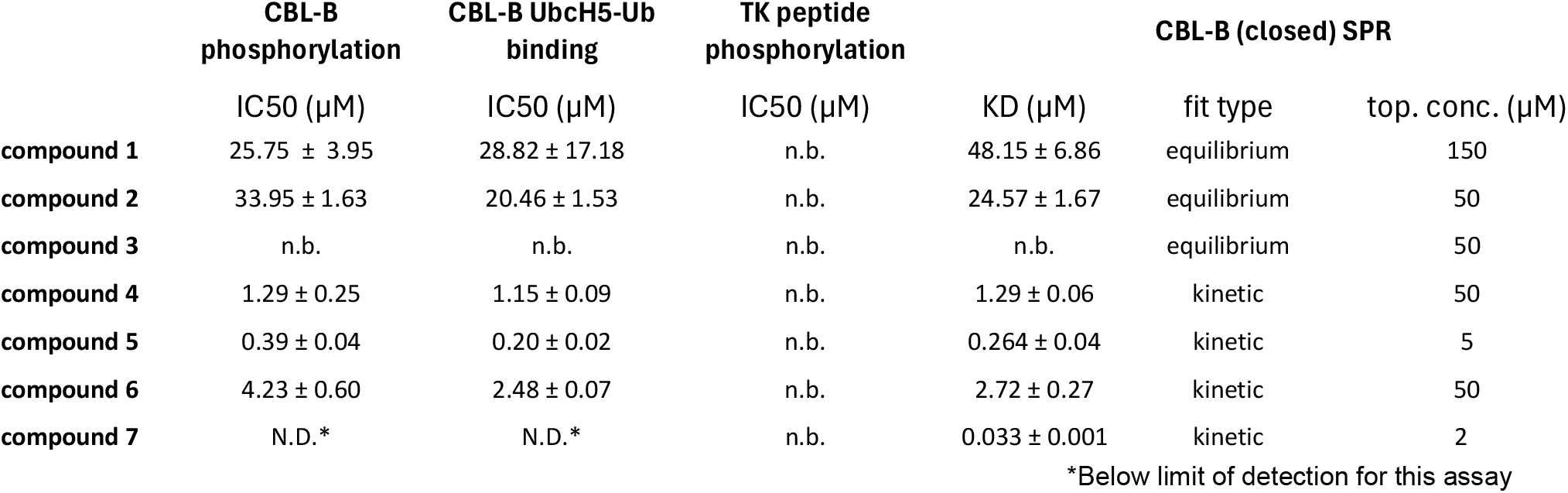
Biochemical triple assay results and biophysical ligand binding to the closed state of CBL-B. N.D. denotes value not determined and n.b. no detectable binding.

|  | CBL-B<br>phosphorylation | CBL-B UbcH5-Ub<br>binding | TK peptide<br>phosphorylation | CBL-B (closed) SPR |  |  |
| --- | --- | --- | --- | --- | --- | --- |
| | IC50 (μM) | IC50 (μM) | IC50 (μM) | KD (μM) | fit type | top. conc. (μM) |
| compound 1 | 25.75 ± 3.95 | 28.82 ± 17.18 | n.b. | 48.15 ± 6.86 | equilibrium | 150 |
| compound 2 | 33.95 ± 1.63 | 20.46 ± 1.53 | n.b. | 24.57 ± 1.67 | equilibrium | 50 |
| compound 3 | n.b. | n.b. | n.b. | n.b. | equilibrium | 50 |
| compound 4 | 1.29 ± 0.25 | 1.15 ± 0.09 | n.b. | 1.29 ± 0.06 | kinetic | 50 |
| compound 5 | 0.39 ± 0.04 | 0.20 ± 0.02 | n.b. | 0.264 ± 0.04 | kinetic | 5 |
| compound 6 | 4.23 ± 0.60 | 2.48 ± 0.07 | n.b. | 2.72 ± 0.27 | kinetic | 50 |
| compound 7 | N.D.* | N.D.* | n.b. | 0.033 ± 0.001 | kinetic | 2 |

\*Below limit of detection for this assay

### Structural Characterization of the Singleton Hit

We separated the enantiomers of compound **1** (Figure 2 A) and tested both isomers in biochemical assays and by SPR (Figure 2 B-C, Table 1). Compound **2** displayed a robust FRET signal in the phosphorylation and E2 binding assays without kinase interference in the control whereas its enantiomer, compound **3** exhibited no appreciable signal in all three biochemical assays. SPR confirmed that compound **2** is the active eutomer, with a K_D_ of 24.57 μM and an approximately two-fold higher affinity than the racemic HTS hit. We prepared CBL-B apo crystals and crystals of a spry-2 peptide bound, phosphorylated CBL-B in complex E2~Ub to enable structural follow up of HTS hits, but neither specimen was suitable to resolve compound **2** via soaking (Figure 3, Supplemental Table 1). We therefore co-crystallized compound **2** with CBL-B and solved the structure to 1.75Å resolution (Figure 2 D, PDB 9XZB) and confirmed the stereochemical assignment of compound **2** as the (S)-isomer.

**Figure 2:**
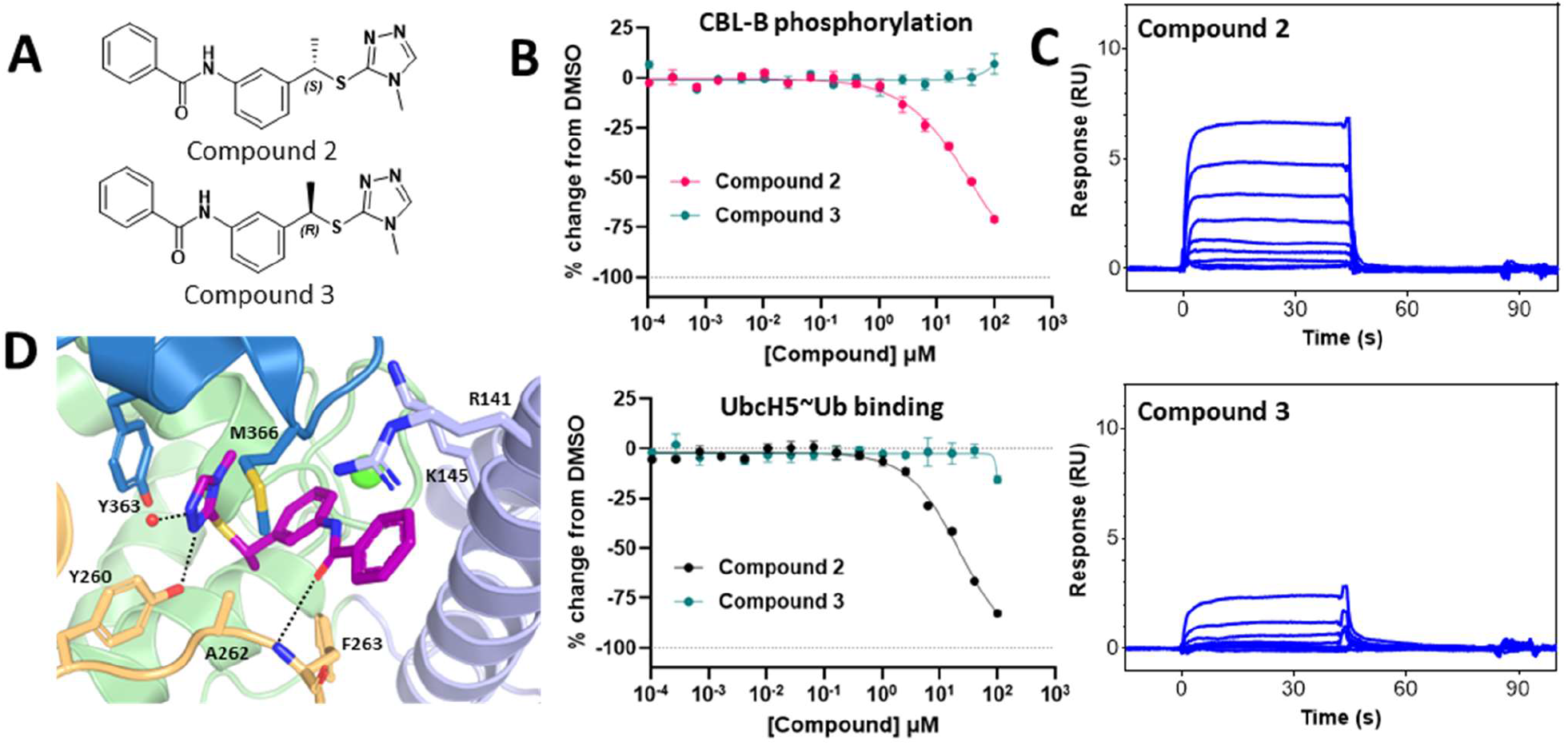
Characterization of the chirally resolved HTS hit. A) Structures of the enantiopure compounds **2** and **3** obtained from chiral separation of the racemic HTS hit, compound **1** B) Biochemical activity of compounds **2** and **3**. Compound **2** was identified in the phosphorylation assay (red data series) and the UbcH5B~Ub binding assay (black data series) as the eutomer with no appreciable binding from the distomer compound **3** (green data series in both) C) Representative SPR sensograms of compound **2** binding to the unphosphorylated closed state of CBL-B with a K_D_ of 24.6 μM. Sensograms for compound **3** could not be fit to a binding model to derive a K_D_. D) Crystal structure of compound **2** bound to CBL-B reveals S-enantiomer as eutomer species. CBL-B is shown as cartoon with 4HB domain (res. 36-168) in pale blue, EF-Hand (res. 169-241) in pale green, SH2 (res. 242-337) in pale orange, and LHR (res. 338-369) in medium blue. A calcium atom is shown as a green sphere, and a small red sphere represents a tightly bound water molecule. The ligand (purple sticks) binds into a shallow cavity between all four domains with prominent sidechains shown as sticks. Hydrogen bonding interactions are shown as dashed black dotted lines.

**Figure 3:**
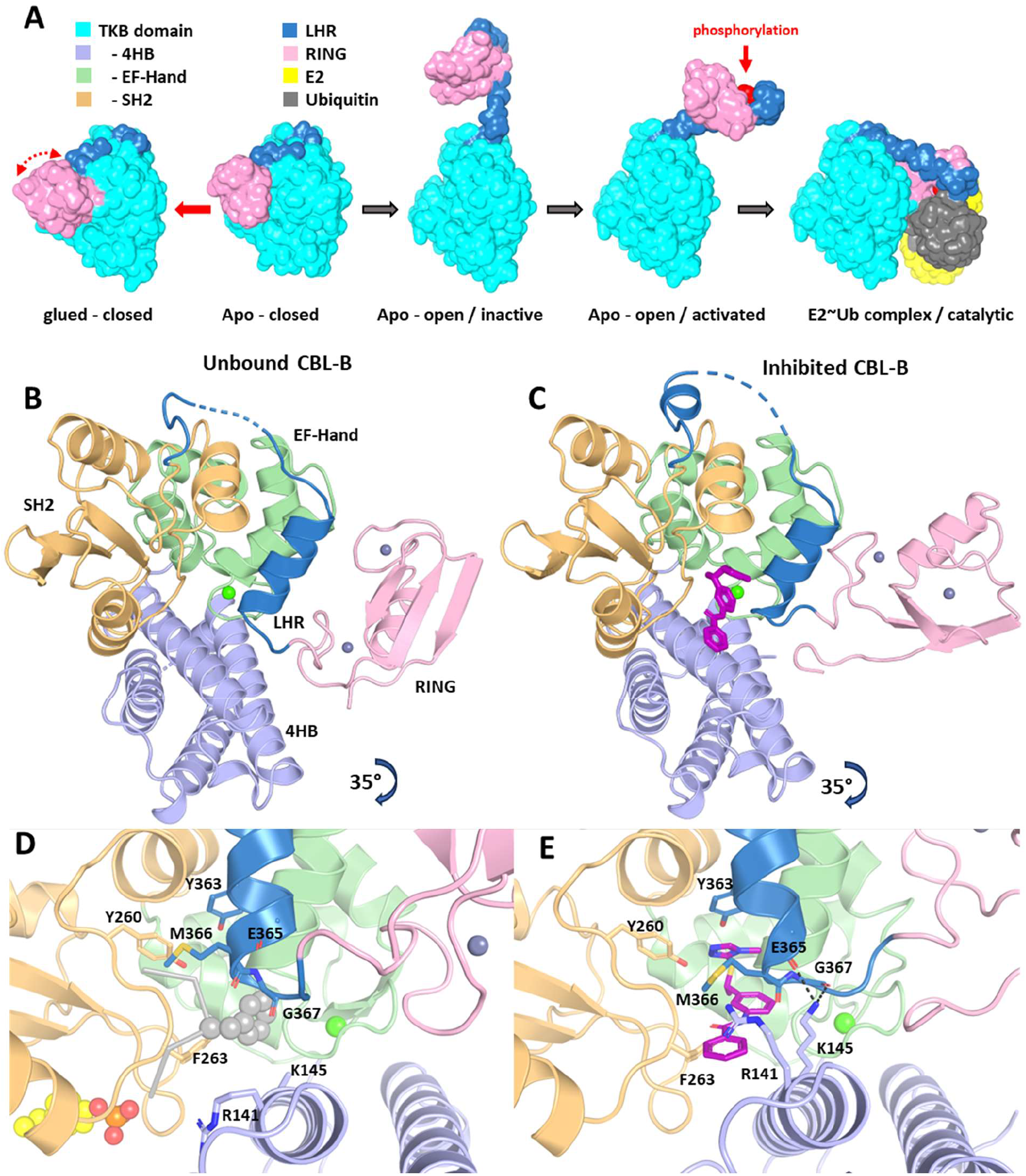
Inhibition mechanism of compound 2. The CBL-B protein structure is colored by subdomains as follows: 4HB (res 36-168) in pale blue, EF-Hand (res 169-241) in pale green, SH2 (res 242-337) in pale orange, LHR (res 338-369) medium blue, and RING (res 370-426) pink. Calcium and zinc atoms are shown as green and grey spheres, respectively. A) Model of the CBL-B activation sequence with the TKB domain (res 36-337, 4HB, EF-Hand, SH2) for simplicity shown in cyan. The substrate peptide binds on the opposite side of the TKB in this view. Upon displacement of the LHR/RING by peptide binding Y363 phosphorylation (red) turns the enzyme active and capable of forming the catalytic complex with E2 (yellow) and Ubiquitin (dark grey). B) Overview of CBL-B apo - closed crystal structure (PDB ID 9XZA) with labeled subdomains. Binding location of compound **2** in a pivotal location between the first four subdomains of CBL-B in the glued - closed state. The RING domain undergoes a significant relocation upon ligand binding under the LHR helix. See text for details. D) Closeup view on the C-terminus of the LHR helix in the CBL-B apo – closed crystal structure. Additional representations are the SH2 loop from the catalytic complex (PDB ID 9ZBH) in light grey ribbons with spheres showing the sidechains of F263 and sprouty 2 phosphotyrosine pY55 (yellow carbon atoms). E) Pi stacking of the triazole with Y363 is the primary sidechain contact of the ligand. M366 sidechain is coordinated centrally over the kinked ligand. Compound **2** binding induces a closure of the gap between the 4HB domain and the LHR helix as R141 migrates over the terminal benzene of the ligand and a niche-3 structure forms between K145 and the carbonyls of E365 and G367.

The ligand binding site is located between the three subdomains of the TKB and the LHR with no direct contact to the RING domain (Figure 3 C). Compound **2** binds in a cavity under the LHR helix with most of the target engagement originating from the 1,2,4-triazole moiety. Key interactions include a π−π stacking interaction with Y363, a hydrogen bond with the phenol of Y260, and a tightly bound water molecule. Additionally, the sidechain of M366 moves to the other side of the triazole to make VDW interactions with the central aromatic ring of the ligand and a subsequent distortion of the following LHR turn into a 3_10_ helical conformation. Upon this LHR transformation, a niche-3 motif at the carbonyls of E365 and G367 forms with the sidechain amine of K145 and a closure of the gap between the 4HB and the LHR by ~1Å (Figure 3 D-E). This forces the peptide chain exiting the LHR closer to the EF-hand and the RING domain, causing it to reposition along the EF-hand closer to the 4HB domain (Figure 3 E). At the SH2 domain, the amide carbonyl of compound **2** forms a hydrogen bond with the mainchain amide nitrogen of F263, blocking its sidechain from displacing the LHR in response to substrate binding during the early CBL-B activation sequence (Figure 3 A & D). These observations suggest that the inhibitor acts as an intramolecular glue that induces de novo contacts among four subdomains of CBL-B. In this stabilized conformation, the central tyrosine (Y363) is sequestered and therefore inaccessible to kinase-mediated activation, while the LHR becomes tethered to the TKB domain. This intramolecular engagement competes with binding of tyrosine-phosphorylated substrate peptides and ultimately results in inhibition of the ligase.

### Hit Optimization and T-cell activation

Initial exploitation of the structure-activity relationship was directed at exploring substitutions for the benzene moiety of the hit compound (Figure 4, Table 1). An isoquinoline substitution (compound **4**) led to a 20-fold increase in affinity (K_D_ = 1.29 ± 0.06 μM) and activity due to a better fill of the cavity between the 4HB and SH2 domains. A trifluoromethyl pyridine substituent at the same position in compound **5** gave an additional five-fold potency increase K_D_ = 0.26 ± 0.04 μM) through improved contacts with the alkyl chains of R141 and K141, without disrupting the niche-3 motif (Figure 4). We also synthesized and characterized the distomer compound **6** that shows a 10-fold lower activity, consistent with unfavorable conformational energy of the chiral methyl group in that ligand binding pose.

**Figure 4:**
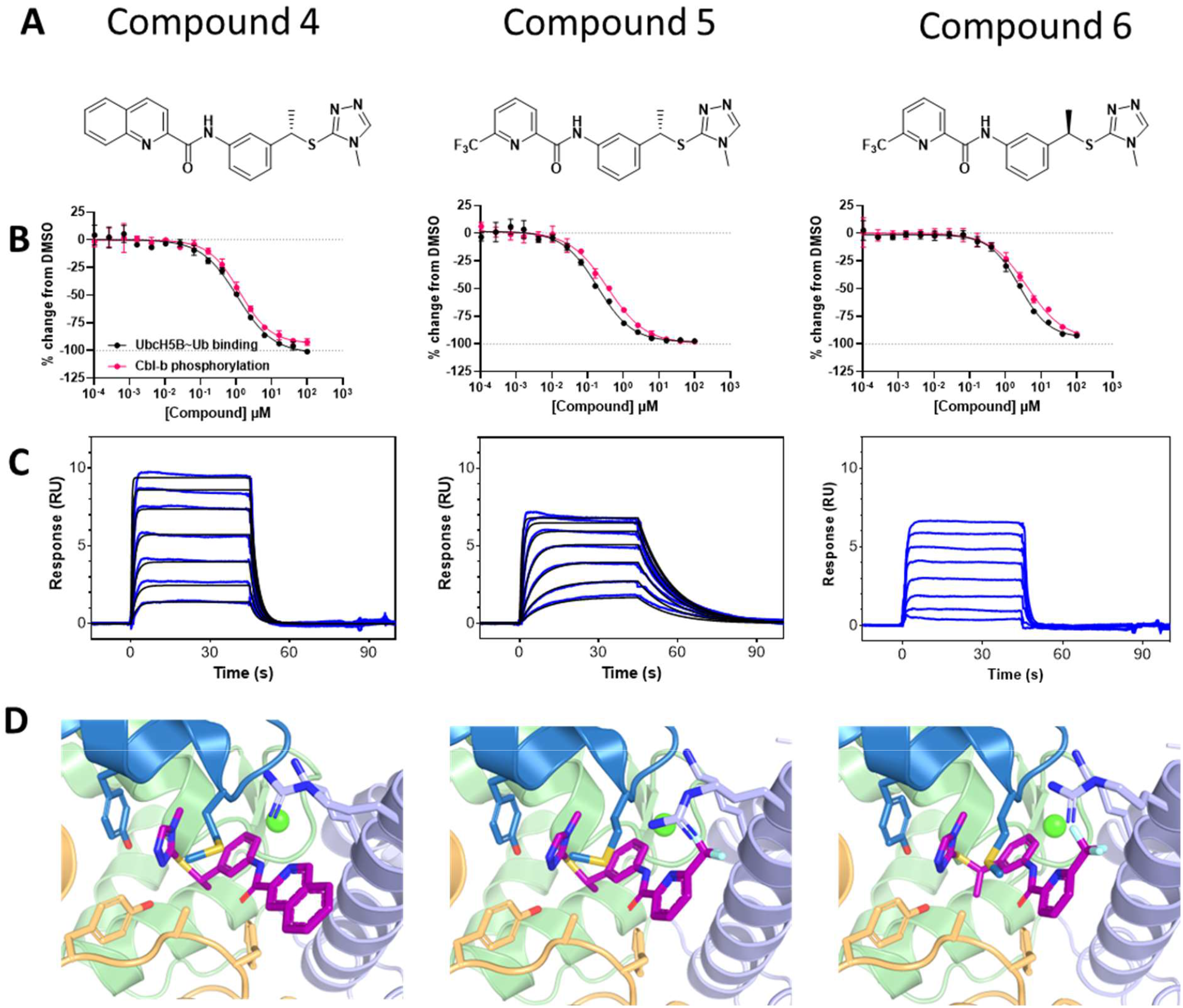
Structure Affinity Relationship of follow-up designs based on the HTS hit. A) Structures of compounds **4-6**. Compounds **4** and **5** were designed to improve space filling between the 4HB and LHR domains relative to compound **2**. Compound **6** is the distomer of compound **5**. B) Activity of compounds **4-6** in the CBL-B phosphorylation and UbcH5B~Ub assays. C) Representative SPR sensograms for compounds **4-6** binding to CBL-B. D) Crystal structures of compounds **4-6** bound to CBL-B (PDB ID: 9XZC, 9XZD, 9XZE). The structures illustrate a more complete fill of the gap between LHR and 4HB under the aliphatic portion of R141 and K145 side chains.

An early functional readout of the lead series was obtained by monitoring human T-cell activation following titration with compound **5**, using the distomer, compound **6**, as a control to assess target specificity. We monitored surface expression of activation markers CD69 and CD25 and secretion levels of IFN-γ and IL-2 cytokines (Figure 5 C-E) from PBMCs under CD3-only or CD3 and CD28 co-stimulation conditions (Figure 5). The early T-cell activation marker CD69 showed a modest increase in surface expression after 48 hours of incubation for both stimulation conditions when treated with compound **5**, but not with compound **6** (Figure 5 A). Similarly, levels of the late T-cell activation marker CD25 increased four-fold in both stimulation conditions for compound **5**, whereas no increase was observed for compound **6** (Figure 5 B). PBMCs treated with compound **5** also showed a modest increase of IFN-γ secretion especially in presence of co-stimulation, despite the late timepoint readout (Figure 5 C). The strongest response elicited by compound **5** treatment was seen in secretion of IL-2, a cytokine that promotes T cell survival, proliferation, and differentiation. At the highest concentration, IL-2 secretion increased five-fold over background levels induced by CD3/CD28 co-stimulated cells, while compound **6** did not induce additional IL-2 secretion under these conditions.

**Figure 5:**
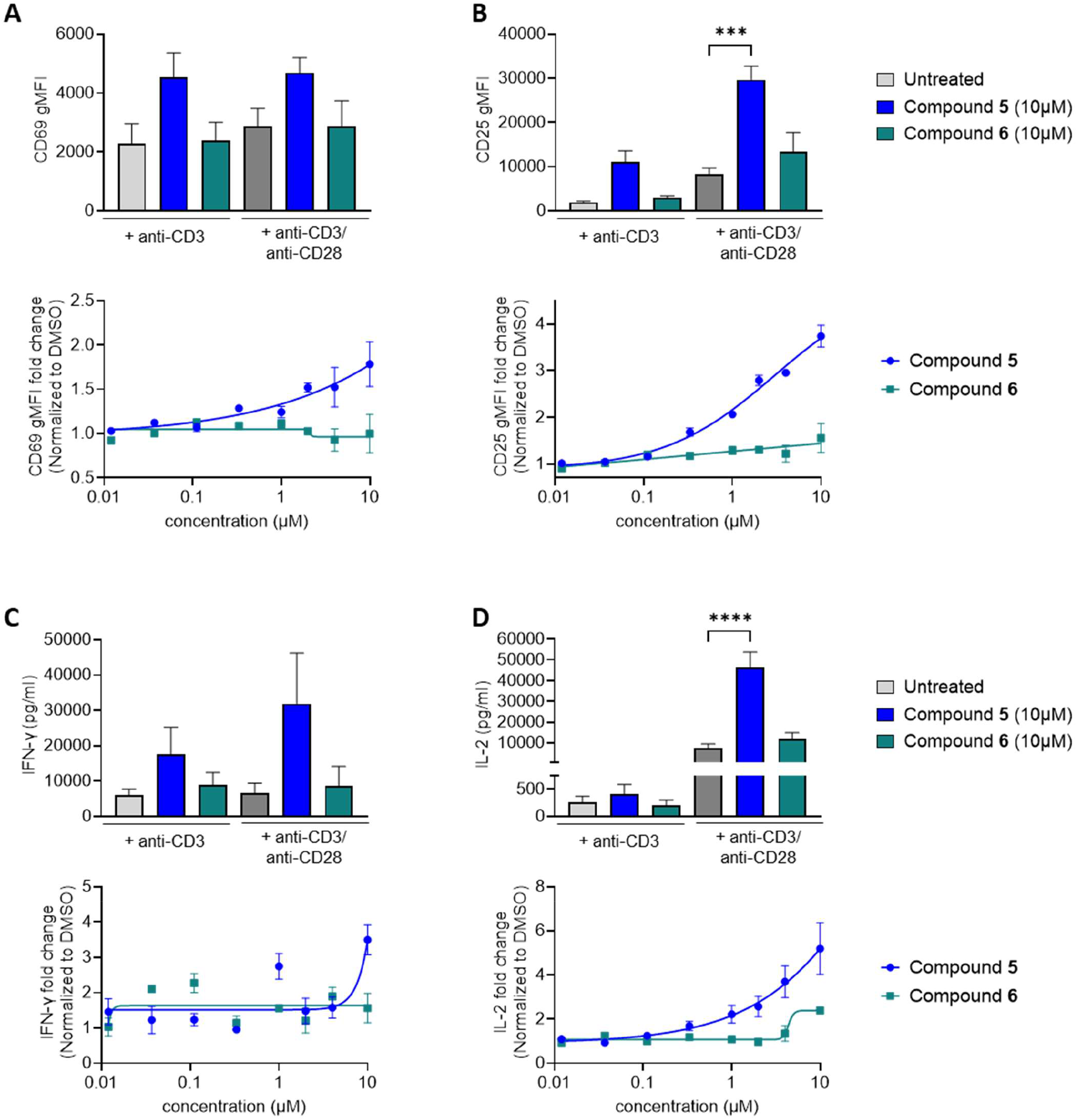
Compound 5 induces activation of T cells. T cells derived from human PBMCs from adult, healthy donors (n=3) were treated with either the enantiomer pair compound **5** or compound **6** for 1 hour prior to stimulation. For each panel, the bar plot shows the effect of the highest concentration, and the line graph represents non-linear fitted curves of the dose response, with mean ± SEM. (A-B) CD69 and CD25 expression were quantified using geometric mean fluorescence intensity (gMFI). Fold increase in CD69 and CD25 expression on cells was calculated by dividing the CD69 gMFI and CD25 gMFI of compound treated cells by DMSO (vehicle) background stimulated cells, for each individual donor. C-D) Concentration or fold change of IFN-y and IL-2 levels in CD3/CD28 co-stimulated cells treated with compound **5** or **6**. Bars represent mean + SEM. Statistical significance between treatment groups was calculated using ordinary one-way ANOVA with Dunnett’s multiple comparisons test. ***, P<0.001, ****, P<0.0001

### Tumor growth suppression

Further optimization in the linker region of the lead compound series, replacing the chiral methyl group with an oxetane, resulted in compound **7**, with a K_D_ of 33 nM (Figure 6, Supplemental Figure 1). We first tested compound **7** in the same primary human T cell *in vitro* cytokine secretion assay, where we observed a concentration-dependent increase in IFN-γ for both CD3 stimulated and CD3/CD28 co-stimulated cells (Figure 6 A). IL-2 secretion showed a remarkable twenty-fold increase in CD3 stimulated cells and fifteen-fold increase in CD3/CD28 co-stimulated cells at 10 μM dose concentration (Figure 6 B). Notably, IL-2 secretion did not plateau across the treatment range in CD3/CD28 co-stimulated cells indicating that a maximum level of IL-2 production was not reached within the tested concentration range (Figure 6 B). As a measure of general compound specificity, **7** was screened against a panel of 55 enzymes, receptors and ion channels at 10 μM. No target was inhibited more than 40%, indicating that 7 is not a promiscuous inhibitor (Supporting Information)

**Figure 6:**
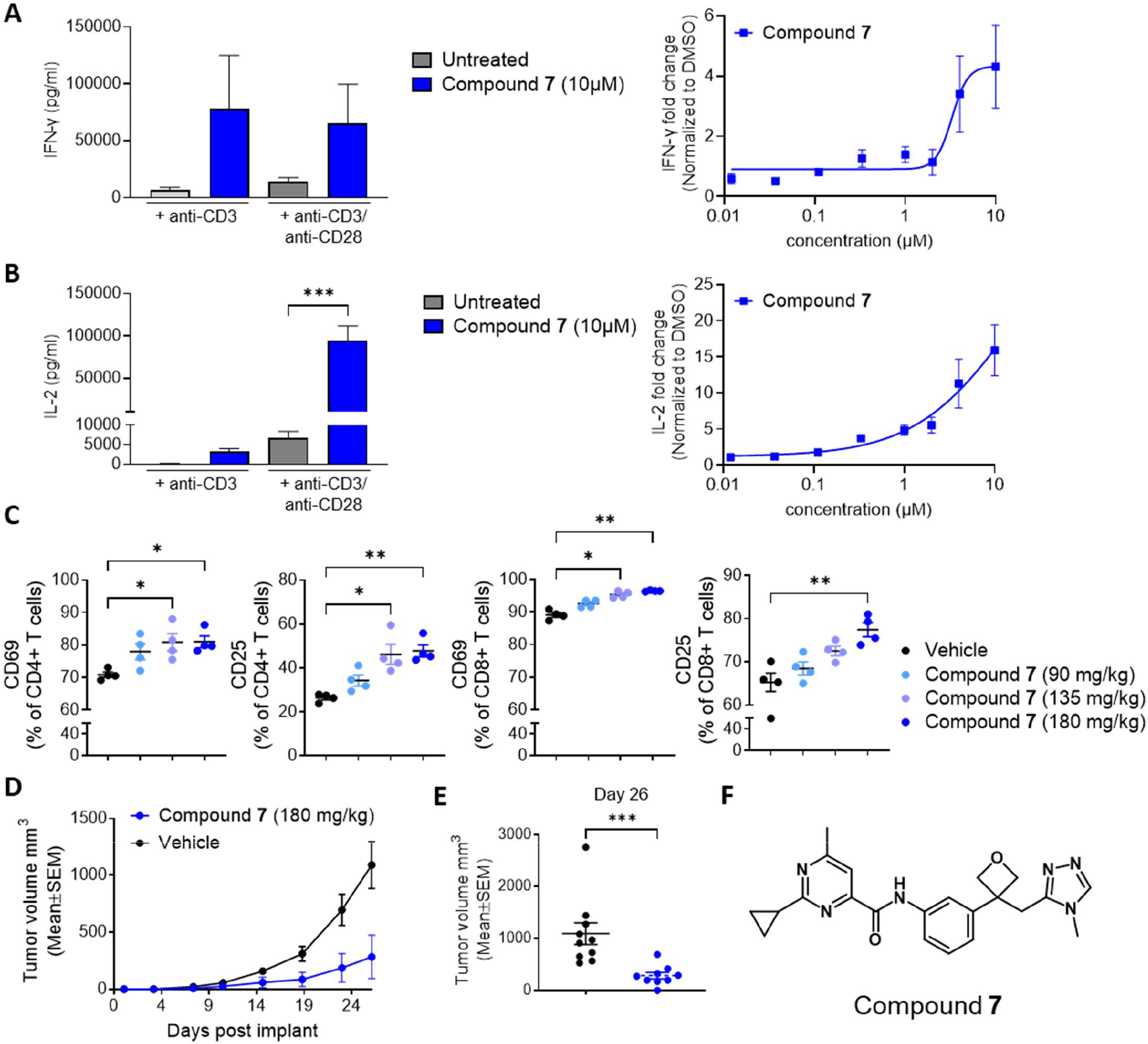
Compound 7 potently enhances T-cell activation and oral administration inhibits tumor growth in a mouse syngeneic tumor model. A-B) Cytokine secretion from compound **7** treated T cells derived from human PBMCs from adult, healthy donors (n = 3). Left panels: bar plots showing IFN-γ (A) and IL-2 (B) secretion in CD3-only and CD3/CD28 co-stimulated cells in the presence or absence of 10 μM compound **7**. Right panels: titration of compound **7** up to 10 μM in CD3/CD28 co-stimulated cells. For each panel, the bar plot shows the effect of the highest concentration, and the line graph represents non-linear fitted curves of the dose response, with mean ± SEM. C) Mice were dosed orally with vehicle or compound **7** (90, 135, or 180 mg/kg) 15 min prior to anti-CD3 antibody injection. Surface expression of CD69 and CD25 on CD4^+^ and CD8^+^ T cells was measured 24 h after antibody injection. D-E) Mice bearing CT26 tumors were treated from Day 3 to Day 26 by oral administration of either vehicle or compound **7** at 180 mg/kg, BID. Beginning day 3, tumor volumes were measured twice weekly. Group mean tumor volumes ± SEM through day 26 are shown in D. Statistical significance of differences in mean tumor volumes between groups was evaluated from days 3 to day 26 using repeated measures two-way ANOVA with Šídák’s multiple comparisons test. Figure 6E shows individual tumor volumes on day 26 with lines at group mean values. Statistical significance of differences in mean tumor volumes between groups on Day 26 was evaluated using Mann-Whitney t-test. *, P<0.05, **, P<0.01, ***, P<0.001. F) Compound **7**.

Compound **7** was used to study the *in vivo* effects of CBL-B inhibition on the expression of T-cell activation markers in the setting of anti-CD3 stimulation. Intravenous administration of anti-CD3 antibody to mice provides antigen independent signaling of the TCR complex, resulting in T-cell activation that can be detected by monitoring expression of activation markers on the surface of T cells (29). Stimulation of mice with anti-CD3 antibody combined with administration of compound **7** at 90, 135, or 180 mg/kg caused a significant dose-dependent increase in the percentage of both CD4^+^ and CD8^+^ T cells expressing the activation markers CD69 and CD25 (Figure 6 C).

The anti-tumor activity of compound **7** as a single agent was evaluated in the murine CT26 tumor model. Compound **7** administered twice daily (BID) at 180 mg/kg resulted in a day 26 mean tumor volume of 283 mm^3^ with individual tumor volumes significantly smaller than vehicle control tumors, corresponding to a tumor growth inhibition of 74% (Figure 6 D-E).

## Discussion

In this study, we report the design and execution of an HTS campaign to identify small molecule inhibitors of CBL-B, a negative regulator of T-cell activation. The screening strategy was designed to exploit a conformational vulnerability of the CBL-B protein in its closed, autoinhibited state when Y363 is buried and inaccessible to phosphorylation. Once phosphorylated by a kinase, CBL-B becomes locked in the activated open conformation unless dephosphorylated. Given that Y363 phosphorylation is required for ligase activity and that steric hindrance precludes its modification in the closed conformation, we hypothesized that stabilizing the autoinhibited state could block CBL-B activation and thereby render T cells more sensitive to activation. We hypothesized that intramolecular glue reinforcing interactions between the LHR helix and TKB domain, and preventing the F263-mediated displacement, could serve as an effective inhibitor of the conformational change that underlies phosphorylation-driven activation. Based on this rationale, we prioritized the CBL-B phosphorylation assay for HTS and incorporated a kinase counter screen to rule out false positives arising from Src kinase inhibition.

The high initial hit rate observed in this E3 ligase closed state screen necessitated the development of a stringent hit validation funnel consisting of dose-response assays for CBL phosphorylation and E2 binding, a Src kinase inhibition counter screen, and orthogonal measurement of biophysical binding, which ultimately provided high confidence for a singleton hit with low affinity. An additional HTS using phosphorylated, CBL-B protein was designed to measure recruitment of ubiquitin-charged E2 (UbcH5B~Ub) in a FRET-based assay to capture compounds that inhibit E2 binding as a complementary mechanism to block CBL-B function. No hits were identified in this screen.

Chiral resolution of the closed state singleton hit revealed a preferred stereoisomer, further supporting a specific interaction with CBL-B. Crystallographic analysis of compound **2** revealed direct engagement with Y363 and F263, effectively obstructing CBL-B activation through F263-mediated LHR displacement, inducing a local deformation in the LHR helix, and a shift in RING domain localization. At the time, this CBL-B conformation had not yet been described. Initial analysis of the structure led us to propose that the niche-3 contact between K141 and the carboxyterminal end of the LHR is a native interaction that stabilizes the closed conformation, firmly enforcing SH2 substrate binding for CBL-B activation. The lead compound is engaging both the LHR and TKB domains and thus stabilizes this autoinhibited state as an intramolecular glue.

Although the initial hit displayed low potency, this structural insight guided early affinity optimization. Expanding the solvent-exposed benzamide yielded a tenfold improvement in activity, and further iterations closing the void volume at the niche-3 pocket beneath K141 produced compound **5**, which demonstrated submicromolar potency and functional activity in T-cell assays. The chirality-dependent T-cell activation observed with compound **5** and its distomer compound **6** confirmed that cytokine secretion and expression of T-cell activation markers were on-target effects resulting from stereoselective engagement of CBL-B. Strikingly, IL-2 secretion did not reach a plateau in the dose range tested, suggesting further potency improvement was needed to achieve a full response. Encouraged by that finding we continued lead-optimization and arrived at compound **7**, which exhibited another tenfold potency increase over **5**. We used this compound to recapitulate the phenotype of a CBL-B point mutation that disrupts E2 binding (8). First, we showed in human CD3+ T-cells that CBL-B uncouples T-cell activation from CD28 co-stimulation, along with a dose-dependent enhancement of IL-2 secretion. Next, we demonstrated a significant dose-dependent increase in cell-surface activation markers *in vivo* in both CD4^+^ and CD8^+^ T cells. Finally, we showed *in vivo* tumor growth inhibition in a syngeneic mouse model by oral administration of compound **7**, confirming the functional relevance of these immunological effects. Further optimization for affinity and pharmacokinetics resulted in NX-1607, the first CBL-B inhibitor to enter clinical testing (clinical trials.gov ID: NCT05107674).

Together, these findings demonstrate that the CBL-B E3 ligase is a druggable target that can be inhibited by a small molecule intramolecular glue which recapitulates the T-cell activation and anti-tumor activity phenotypes observed with a CBL-B genetic knockout. The results suggest that rational screening strategies designed to identify inhibitors targeting an autoinhibited protein conformation are a technically feasible approach for drugging E3 ligases, indicating such strategies may be more broadly applicable than appreciated, particularly for multi-domain proteins that assume multiple conformational states.

## Materials and Methods

### Protein production

The coding sequence for CBL-B residues 36-427 was inserted in a pGEX4T1 expression vector to yield a N-terminal GST fusion protein with thrombin cleavage site prior to the CBL-B sequence. For streptavidin conjugation, a construct was made with an Avi-tag sequence insertion between the thrombin cleavage site and CBL-B. The protein was expressed in BL21 DE3 cells growing exponentially in TB medium. Temperature was reduced to 18 °C and induced at OD 600 1.7 with 0.5 mM IPTG overnight. Cells were pelleted 25 min at 3000 rpm and resuspended in 500 ml Lysis buffer (200 mM NaCl, 50 mM Tris-HCl pH 7.5, 2 mM DTT, 1x Protease inhibitor cocktail). Cells were lysed twice with 15,000 PSI in a microfluidizer and Lysate centrifuged 1 hour at 13,000xg. Supernatant was centrifuged 30 min at 30,000xg and filtered with 0.45 um filter. 12ml GST beads were added to the cleared lysate and incubated 1 hour at 4 °C and 75 rpm. Sample was poured over a glass column and beads washed with 300 ml Lysis buffer without protease inhibitor and then with 50 ml overnight cleavage buffer (200 mM NaCl, 50 mM Tris-HCL pH 8.0, 2.5 mM CaCl_2_ and 2 mM DTT). Column outlet was closed and beads resuspended in 13 ml overnight cleavage buffer supplemented with 0.1 mg/ml (final) thrombin and incubated overnight. Next morning the sample was captured in a 50 ml Falcon tube and remaining protein was washed out with 45 ml buffer. The sample was diluted to 250 ml using 50 mM Tris HCl pH 8.0 2 mM DTT (buffer A) and then applied to a 5 ml Q Sepahrose HP column equilibrated in buffer A and washed with 20% Buffer B (1000 mM NaCl, 50 mM Tris-HCl pH 8.0, 2mM DTT) and eluted in a gradient from 20% to 60% buffer B over 10 CV. Peak fraction was captured in 50 ml Faclon tubes, concentrated to 2 ml and run over a 16/600 Superdex 75 size exclusion column equilibrated in 200 mM NaCl, 20 mM Tris-HCL pH 8.0, and 2 mM DTT.

### CBL-B phosphorylation assay

Phosphorylation assays were performed in a 384-well plate at room temperature in a 10 µl reaction volume by pre-incubating 150 nM biotinylated CBL-B (residues 36-427) in an assay buffer containing 50 mM HEPES pH 7.5, 50 mM NaCl, 0.01% Triton X-100, 0.01% BSA and 1 mM DTT in the presence of a candidate compound in 1% DMSO (final concentration) for 1 hour. After incubation in the presence of the candidate compound, the plate was incubated with the Src kinase mix (2 nM Src kinase (residues 248-533) and ZAP70 peptide, 5 mM ATP and 25 mM MgCl_2_). After 1 hour incubation with the kinase, the reactions were stopped with 450 mM EDTA and incubated for an additional 1 hour with TR-FRET with 25 nM Streptavidin-XL-665 and 0.25 x ptyrosine-Antibody-Terbium (Revvity) for phospho-tyrosine detection. Fluorescence signals were measured at two wavelengths, 665 nm, and 485 nm, following excitation at 340 nm using an Envision plate reader (Perkin Elmer). TR-FRET signal was calculated using a 665/485 nm ratio. Data were corrected to subtract background signal in the absence of Src kinase. Inhibitor activity was calculated as percent change in phosphorylation compared to the signal from 150 nM fully phosphorylated CBL-B. A standard Hill equation (Y = Bmin + ((Bmax − Bmin) /(1+(IC_50_/X)^Hill^))) was fit to the dose response data in ActivityBase (IDBS) to obtain the IC_50_ values.

### Peptide phosphorylation assay

Phosphorylation assays were performed in a 384-well plate at room temperature in a 10 µl reaction volume by pre-incubating 3 nM BodipyTMR labeled TK peptide in an assay buffer containing 50 mM HEPES pH 7.5, 50 mM NaCl, 0.01% Triton X-100, 0.01% BSA and 1 mM DTT in the presence of a candidate compound in 1% DMSO (final concentration) for 1 hour. After incubation in the presence of the candidate compound, the plate was incubated with the Src kinase mix (2 nM Src kinase (residues 248-533) and ZAP70 peptide, 5 mM ATP and 25 mM MgCl_2_). After 1 hour incubation with the kinase. the reactions were stopped with 300 mM EDTA and incubated for an additional 1 hour with TR-FRET phospho-tyrosine detection reagents (0.25 x pTyrosine-Antibody-Terbium). Fluorescence signals were measured at two wavelengths, 570 nm, and 485 nm, following excitation at 340 nm using an Envision plate reader (Perkin Elmer). TR-FRET signal was calculated using a 570/485 nm ratio. Data were corrected to subtract background signal in the absence of Src kinase. Inhibitor activity was calculated as percent change in phosphorylation compared to the signal from 3 nM fully phosphorylated TK peptide. A standard Hill equation (Y = Bmin + ((Bmax − Bmin) /(1+(IC50/X)^Hill^))) was fit to the dose response data in ActivityBase (IDBS) to obtain the IC_50_ values.

### CBL-B E2-Ub binding assay

Phosphorylation assays were performed in a 384-well plate at room temperature in a 10 µl reaction volume by pre-incubating 12 nM biotinylated CBL-B (residues 36-427) in an assay buffer containing 50 mM HEPES pH 7.5, 100 mM NaCl, 0.01% Triton X-100, 0.01% BSA and 1 mM DTT in the presence of a candidate compound in 1% DMSO (final concentration) for 1 hour. After incubation in the presence of the candidate compound, the plate was incubated with the Src kinase mix (60 nM Src kinase (residues 248-533) and ZAP70 peptide, 5 mM ATP and 25 mM MgCl_2_). After 1.5-hour incubation with the kinase the reactions were stopped with 300 mM EDTA and incubated for an additional 1 hour with TR-FRET orthogonal detection reagents (2.5 nM Streptavidin-Terbium and 250 nM UbcH5b~Ub-BodipyFL). Fluorescence signals were measured at two wavelengths, 520 nm, and 620 nm, following excitation at 340 nm using an Envision plate reader (Perkin Elmer). TR-FRET signal was calculated using a 520/620 nm ratio. Data were corrected to subtract background signal in the absence of Src kinase. Inhibitor activity was calculated as percent change in phosphorylation compared to the signal from fully phosphorylated CBL-B. A standard Hill equation (Y = Bmin + ((Bmax − Bmin) /(1+(IC_50_/X)^Hill^))) was fit to the dose response data in ActivityBase (IDBS) to obtain the IC_50_ values.

### SPR

SPR data were collected using Cytiva T200 and S200 biosensors (Cytiva Life Sciences) using Cytiva Series-S CM5 sensor chips. Chips were first prepared by immobilizing approximately 10,000 resonance units (RUs) of NeutrAvidin (Invitrogen) to all four flow cells via amine coupling chemistry at 25 °C in an immobilization buffer of 50 mM HEPES, pH 8.0, 150 mM NaCl, and 0.005% Tween-20. The surfaces were treated with a 7-minute injection of 195.5 mM EDC/50 mM NHS followed by a 150-second injection of 125 µg/mL NeutrAvidin diluted in a preconcentration buffer of 50 mM sodium acetate, pH 4.4. Surfaces were then exposed to a three-minute injection of 0.5 M ethanolamine, pH 8.0. Next, 0.2 µM N-terminal Avi-tagged biotinylated CBL-B (36-427) was captured on flow cells 2, 3, and 4 to a density of approximately 1600 RUs. Flow cell 1 served as a reference surface and was prepared without Cbl protein. Finally, 3 mM EZ Link Amine PEG2 Biotin (Thermo Scientific) was flowed over entire chip in two 30 second injections at 30 µL/min to block remaining biotin binding sites. CBL-B immobilization and binding measurements were carried out in assay buffer containing 50 mM HEPES, pH 7.5, 100 mM NaCl, 1 mM TCEP, 0.005% Tween-20, and 2% DMSO at 25 °C. Compounds with uM affinities were serially diluted 2-fold, for 9 points in assay buffer from a top concentration of 50 µM, while those with nM affinities were prepared at a top concentration of 5 µM and the same dilution scheme. Samples were injected for 45 seconds at 100 µL/min flow rate and allowed a dissociation time of 60 seconds. The resulting data were DMSO solvent corrected and double referenced prior to analysis. All data were fit to either a 1:1 equilibrium or kinetic binding model, depending on affinity, using Scrubber V2.0c (BioLogic Software), and sensorgrams and fits were exported to GraphPad Prism to prepare final figures. o

### Crystallography

CBL-B protein (residues 36-427) was formulated at 150 uM in 20 mM KPO_4_, 150 mM NaCl and 2 mM DTT and crystallized at 19°C in sitting drop vapor diffusion setting by mixing with an equal volume well solution of 40 mM MOPS, 60 mM HEPES-Na, 30 mM MgCl_2_, 30 mM CaCl_2_, 8.4% PEG 8000, 15% ethylene glycol in 96-well MCR 2 plates. The wells were filled with 70 ul well solution and covered with 15 ul Al′s Oil (Hampton) to slow down vapor diffusion. Seed stock from apo CBL-B crystals were used to discover ligand co-crystallization conditions. The active CBL-B complex was prepared as described previously (26) but with a sprouty-2 peptide (IRNTNE{PTR}TEGPT) and crystallized in 9.5% PEG 3350, 100 mM sodium formate, 100mM BICINE pH 9.0. Ligand bound CBL-B protein was crystallized with 10-fold ligand excess at 19 °C in sitting drop vapor diffusion setting by mixing with an equal volume well solution of 100 mM MES pH 5.6-6.2, 200 mM LiSO_4_, 16-20% PEG 3350, 50 mM DTT. Rod shaped crystals appeared in 1-3 days. All crystals were cryo protected in their respective well solution adjusted to 20% ethylene glycol and flash frozen in liquid nitrogen for data collection at ALS beamlines 5.0.2. and 8.2.1 and SSRL beamline 12-2. Data frames were processed with XDS (30) and scaled with aimless(31). The structures were solved with Phaser (32) and refined using phenix.refine (33) and coot (34).

### Chemistry

Compound 1 was purchased from Enamine. The synthesis of compounds 2–7 is described in WO2019148005 A1.

### T-cell activation assay

T cells isolated from human peripheral blood mononuclear cells (PBMCs) were seeded in 96-well round bottom plates and treated with the compounds (10 µM, 4 µM, 2 µM, 1 µM, 0.3 µM, 0.1 µM, 0.037 µM and 0.012 µM) for 1 hour at 37°C, followed by transfer to anti-CD3 (clone OKT3, 10 µg/ml) pre-coated 96-well round bottom plates. For co-stimulation, anti-CD28 (clone CD28.2, 5 µg/ml) was added to the wells. Plates were incubated at 37°C for 48 hours. After incubation, cell supernatants were harvested for secreted cytokine analysis by Meso Scale Discovery (MSD). Cells were stained with live/dead dye (ThermoFisher, catalog L23101), followed by staining with antibodies to cell surface markers (CD4 AlexaFluor700; clone OKT4, CD69 eFluor450; clone FN50, CD25 BV605; clone BC96, and CD8 PE-Cy7; clone RPA-T8). Flow cytometry was performed on Attune NxT Acoustic Focusing Flow Cytometer, and data were analyzed using FlowJo and GraphPad Prism. Single, live cells (pan T cells) were gated, and expression of CD69 and CD25 on cells was determined using geometric mean fluorescence intensity (gMFI). Levels of secreted IFN-y and IL-2 were determined by V-Plex Proinflammatory Panel 1 Human kit from MSD following the manufacturer’s protocol. Fold increase in CD69 and CD25 expression on cells was calculated by dividing the CD69 gMFI and CD25 gMFI on cells in compound-stimulated cells by DMSO-stimulated cells, for each individual donor. Fold change in IFN-y and IL-2 levels was calculated by dividing the pg/ml value of cytokines in compound-stimulated cells by DMSO-stimulated cells, for each individual donor.

### *In vivo* mouse studies

To characterize the pharmacodynamic effects of *in vivo* administered compound **7**, mice were dosed orally with either vehicle (25% PEG 300 + 5% Solutol, 70% water) or compound **7** at 90, 135, or 180 mg/kg BID. Fifteen minutes after the first dose, either 2 µg of anti-CD3 antibody or PBS for unstimulated controls was injected iv. Twenty-four hours after study initiation, animals were euthanized, splenocytes were prepared, and the percentage of conventional CD4+ T cells (CD45+ CD3+ CD4+ FoxP3-) or CD8+ T cells (CD45+ CD3+ CD8+) positively expressing the activation markers CD25, CD69 were measured by flow cytometry and compared between compound **7** treated groups and controls.

To assess the antitumor efficacy of compound **7**, tumors were initiated in female BALB/c mice by subcutaneously implanting 3 x 10E5 CT26 murine colon carcinoma cells in one flank. Two days after tumor implantation, study day 3, mice were evenly distributed accordingly to body weight into two groups (10 animals per group). Treatments were given from day 3 to day 26 by administration of either: vehicle (0.5% methylcellulose, 0.2% Tween 80 in deionized water) BID, orally (po) or compound **7** at 180 mg/kg BID, po. Tumor volumes and body weight were monitored twice weekly until the end of the study.

## Abbreviations

BID: Twice a day
CBL-B: Casitas B-lineage lymphoma proto-oncogene b
FRET: Fluorescence resonance energy transfer
HTRF: Homogenous time-resolved fluorescence
HTS: High throughput screen
LHR: Linker helix region
PBMC: Peripheral blood mononuclear cells
RTK: Receptor tyrosine kinase
SH2: Src homology 2 domain
SPR: surface plasmon resonance
TCR: T-cell receptor
TKB: Tyrosine kinase binding

