## Supplemental Information for "Discovery and characterization of small molecule inhibitors of CBL-B that act as intramolecular glue to enhance T-cell anti-tumor activity"

**Supplemental Table 1**: Data collection and refinement statistics for CBLB crystal structures. Values in parentheses indicate the highest resolution shell. N/A indicates that this property is not applicable for this dataset.

|  | Apo | Compound 2 | Compound 4 | Compound 5 | | Compound 6 | Compound 7 | Spry 2 complex |
| --- | --- | --- | --- | --- | --- | --- | --- | --- |
| **PDB** | 9XZA | 9XZB | 9XZC | 9XZD | | 9XZE | 9ZBG | 9ZBH |
| **Experiment** |  |  |  |  | |  |  |  |
| Wavelength [Å] | 0.9795 | 1.0 | 1.0 | 1.0 | | 1.0 | 1.0 | 1.0 |
| Space Group | P 3_1_ | P 2_1_ 2_1_ 2_1_ | C 2 | P 2_1_ 2_1_ 2_1_ | | P 2_1_ 2_1_ 2_1_ | P 2_1_ 2_1_ 2_1_ | P2_1_ |
| Unit Cell |  |  |  |  | |  |  |  |
| (a, b, c,) [Å] | 101.8 101.8 104.6 | 74.7 97.8 57.5 | 98.0 50.0 86.0 | 50.0 71.2 116.0 | | 49.5 69.9 111.2 | 50.0 70.5 114.0 | 95.5 132.2 122.1 |
| (α, β, γ) [°] | 90 90 120 | 90 90 90 | 90 99.2 90 | 90 90 90 | | 90 90 90 | 90 90 90 | 90 92.0 90 |
| **Data** |  |  |  |  | |  |  |  |
| Resolution range [Å] | 15.00-2.30 (2.36-2.30) | 50.00-1.75 (1.80-1.75) | 50.00-1.80 (1.85-1.8) | 50.00-1.65 (1.69-1.65) | 50.00-2.00 (2.05-2.00) | | 50.00-2.00 (2.05-2.00) | 50.00-2.35 (2.41-2.35) |
| Unique Reflections | 51,530 (5,276) | 43,167 (4,215) | 38,161 (3,770) | 49,955 (4,891) | | 26,772 (2,600) | 27,804 (2000) | 125,664 (9223) |
| Completeness [%] | 95.70 (97.69) | 99.87 (98.99) | 99.73 (99.05) | 98.61 (97.47) | | 99.90 (99.54) | 99.5 (99.5) | 99.8 (99.8) |
| Multiplicity | 2.7 (2.7) | 9.1 (6.5) | 5.5 (5.0) | 8.1 (8.5) | | 8.1 (7.9) | 13.3 (13.9) | 3.5 (3.5) |
| I/σ | 9.90 (1.02) | 10.52 (0.93) | 11.22 (0.95) | 13.06 (1.18) | | 12.32 (1.04) | 12.49 (1.23) | 9.3 (0.99) |
| CC(1/2) | 0.997 (0.295) | 0.998 (0.401) | 0.998 (0.367) | 0.999 (0.442) | | 0.999 (0.378) | 0.999 (0.445) | 99.6 (35.7) |
| R-meas | 0.1041 (1.414) | 0.1348 (1.828) | 0.1075 (1.701) | 0.117 (1.968) | | 0.1338 (2.105) | 0.126 (1.937) | 0.129 (1.617) |
| R-pim | 0.062 (0.846) | 0.044 (0.692) | 0.045 (0.736) | 0.041 (0.663) | | 0.047 (0.738) | 0.045 (0.683) | 0.069 (0.915) |
| Wilson B-factor [Å^2^] | 35.31 | 25.94 | 32.03 | 23.12 | | 36.76 | 35.03 | 45.12 |
| **Refinement** |  |  |  |  | |  |  |  |
| Resolution range [Å] | 14.73-2.30 (2.34-2.30) | 48.9-1.75 (1.79-1.75) | 45.21-1.80 (1.84-1.80) | 44.99-1.65 (1.69-1.65) | | 45.24-2.0 (2.05-2.0) | 44.33-2.0 (2.05-2.0) | 49.29 - 2.35 (2.38 - 2.35) |
| Reflections used | 51,504 | 43,155 | 38,151 | 49,956 | | 26,762 | 27800 | 125646 |
| Test set size [%] | 5.1 (5.4) | 4.6 (4.6) | 5.0 (5.0) | 4.0 (4.0) | | 7.5 (7.5) | 7.2 (7.7) | 5.1 (5.3) |
| Twinn operator | -k,-h,-l | N/A | N/A | N/A | | N/A | N/A | N/A |
| Twinn Fraction | 0.45 | N/A | N/A | N/A | | N/A | N/A | N/A |
| R-work | 0.178 (0.289) | 0.196 (0.351) | 0.198 (0.331) | 0.197 (0.381) | | 0.197 (0.312) | 0.210 (0.312) | 0.199 (0.322) |
| R-free | 0.229 (0.352) | 0.228 (0.367) | 0.227 (0.326) | 0.222 (0.374) | | 0.244 (0.348) | 0.261 (0.319) | 0.254 (0.370) |
| Model Atoms | 8,672 | 3,292 | 3,151 | 3,288 | | 3,140 | 3,108 | 19,577 |
| Protein | 8,636 | 3,059 | 2,960 | 3,009 | | 3,005 | 3,000 | 19,337 |
| NRX- Ligand | N/A | 24 | 28 | 28 | | 28 | 30 | N/A |
| Other atoms | 19 | 3 | 23 | 24 | | 21 | 14 | 24 |
| Water | 17 | 206 | 140 | 227 | | 86 | 64 | 216 |
| B-factors, mean [Å^2^] | 50.3 | 31.7 | 45.7 | 31.2 | | 42.9 | 48.0 | 50.7 |
| Protein | 50.4 | 31.5 | 46.0 | 31.1 | | 42.9 | 48.3 | 50.8 |
| NRX- Ligand | N/A | 28.0 | 37.5 | 21.2 | | 37.7 | 36.7 | N/A |
| Other atoms | 54.1 | 27.4 | 49.8 | 39.6 | | 53.9 | 55.1 | 49.4 |
| Water | 35.9 | 34.9 | 40.2 | 33.0 | | 40.9 | 37.7 | 41.2 |
| **Validation** |  |  |  |  | |  |  |  |
| Ramachandran [%] |  |  |  |  | |  |  |  |
| Favored | 96.03 | 97.88 | 97.87 | 98.67 | | 97.91 | 96.33 | 96.90 |
| Allowed | 3.97 | 2.12 | 2.13 | 1.33 | | 2.09 | 2.89 | 2.97 |
| Outliers | 0 | 0 | 0 | 0 | | 0 | 0.79 | 0.12 |
| Rotamer Outliers [%] | 2.06 | 0.61 | 0.33 | 0.32 | | 0.32 | 0.97 | 2.15 |
| All atom clash score | 4.89 | 0.83 | 5.16 | 1.68 | | 6.11 | 10.05 | 5.67 |
| Deviation from Ideality |  |  |  |  | |  |  |  |
| Bonds | 0.002 | 0.003 | 0.011 | 0.013 | | 0.013 | 0.009 | 0.008 |
| Angles | 0.499 | 0.572 | 0.998 | 1.195 | | 1.242 | 0.869 | 0.976 |
| Chirality | 0.039 | 0.043 | 0.065 | 0.071 | | 0.064 | 0.050 | 0.054 |
| Planarity | 0.004 | 0.004 | 0.008 | 0.012 | | 0.011 | 0.007 | 0.010 |

**Off-target binding evaluation**

Compound 7 was tested for binding to 55 receptors, ion channels, and transporters at 10 mM via radioligand displacement using scintillation counting.

The results are expressed as a percent of control specific binding:

(measured specific binding / control specific binding) *100

and as a percent inhibition of control specific binding:

100-((measured specific binding / control specific binding) *100)

obtained in the presence of the test compounds. The IC50 values (concentration causing a half-maximal inhibition of control specific binding) and Hill coefficients (nH) were determined by non-linear regression analysis of the competition curves generated with mean replicate values using Hill equation curve fitting

Y=D+[(A-D) / (1+(C/C_50_))^nH^]

where Y = specific binding, A = left asymptote of the curve, D = right asymptote of the curve, C = compound concentration, C50 = IC_50_, and nH = slope factor. This analysis was performed using software developed at Cerep (Hill software) and validated by comparison with data generated by the commercial software SigmaPlot® 4.0 for Windows® (© 1997 by SPSS Inc.). The inhibition constants (Ki ) were calculated using the Cheng Prusoff equation:

Ki= IC_50_ / (1+L/KD)

where L = concentration of radioligand in the assay, and KD = affinity of the radioligand for the receptor. A scatchard plot is used to determine the KD.

**Supplemental Table 2:** Screen results. As a measure of general compound specificity, compound **7** was screened against a panel of 55 receptors, ion channels, and transporters at 10 μM. No target was inhibited more than 40%, indicating that compound **7** is not a promiscuous inhibitor

| Target | 1^st^ Inhibition  [%] | 2^nd^ Inhibition  [%] | Mean Inhibition  [%] |
| --- | --- | --- | --- |
| **Receptors** |  |  |  |
| A1 | 34.3 | 44.4 | 39.3 |
| A2A | -27.4 | 2.3 | -12.5 |
| A3 | -1.0 | -3.2 | -2.1 |
| α1 | -16.1 | 19.3 | 1.6 |
| α2 | -2.3 | 0.4 | -1.0 |
| β1 | 1.9 | 12.0 | 7.0 |
| β2 | 2.1 | 7.7 | 4.9 |
| AT1 | -3.9 | -12.3 | -8.1 |
| BZD | -6.2 | -5.5 | -5.9 |
| B2 | -7.9 | -4.0 | -6.0 |
| CB1 | 1.5 | -0.3 | 0.6 |
| CCK1 (CCKA) | 1.8 | -8.8 | -3.5 |
| D1 | 1.9 | 18.2 | 10.0 |
| D2S | -13.2 | 21.9 | 4.3 |
| ETA | -9.4 | 19.8 | 5.2 |
| GABA | -27.1 | 12.3 | -7.4 |
| GAL2 | -7.1 | 2.5 | -2.3 |
| CXCR2 (IL-8B) | -5.0 | 4.7 | -0.1 |
| CCR1 | -12.5 | -14.8 | -13.7 |
| H1 | -8.1 | -7.6 | -7.9 |
| H2 | 11.3 | 14.0 | 12.6 |
| MC4 | 4.1 | 8.2 | 6.2 |
| MT1 (ML1A) | 2.6 | 5.9 | 4.2 |
| M1 | -11.2 | 7.5 | -1.8 |
| M2 | 14.9 | 25.6 | 20.3 |
| M3 | -7.8 | -1.3 | -4.6 |
| NK2 | -1.7 | 2.7 | 0.5 |
| NK3 | 1.7 | -3.8 | -1.0 |
| Y1 | -11.9 | -3.9 | -7.9 |
| Y2 | -15.3 | -2.6 | -8.9 |
| NTS1 (NT1) | -10.2 | -11.2 | -10.7 |
| δ (DOP) | 6.5 | 7.3 | 6.9 |
| κ (KOP) | -10.9 | -9.2 | -10.1 |
| μ (MOP) | -39.7 | 14.8 | -12.5 |
| NOP (ORL1) | -16.2 | -8.6 | -12.4 |
| EP4 | 17.1 | 10.2 | 13.6 |
| 5-HT1A | 11.4 | 4.9 | 8.1 |
| 5-HT1B | -0.9 | -27.7 | -14.3 |
| 5-HT2A | -12.9 | -1.7 | -7.3 |
| 5-HT2B | -10.7 | -4.1 | -7.4 |
| 5-HT3 | -25.4 | -7.4 | -16.4 |
| 5-HT5a | -11.4 | -13.1 | -12.2 |
| 5-HT6 | 8.6 | 4.3 | 6.4 |
| 5-HT7 | -9.2 | -13.6 | -11.4 |
| sst | -7.6 | -1.7 | -4.6 |
| VPAC1 (VIP1) | -18.2 | -21.8 | -20.0 |
| V1a | 2.3 | 3.3 | 2.8 |
| **Ion Chanels** |  |  |  |
| Ca2+ | 0.4 | 9.4 | 4.9 |
| KV | -5.8 | -0.8 | -3.3 |
| SKCa | 12.4 | 9.8 | 11.1 |
| Na+ | -9.1 | 5.7 | -1.7 |
| Cl- | 5.3 | 8.2 | 6.8 |
| **Transporters** |  |  |  |
| norepinephrine transporter | -27.7 | -23.3 | -25.5 |
| dopamine transporter | 2.3 | 1.9 | 2.1 |
| 5-HT transporter | -4.9 | -4.8 | -4.9 |

Biophysical characterization of Compound 7.

Supplemental Figure 1: Compound 7 SPR sensograms and crystal structure.

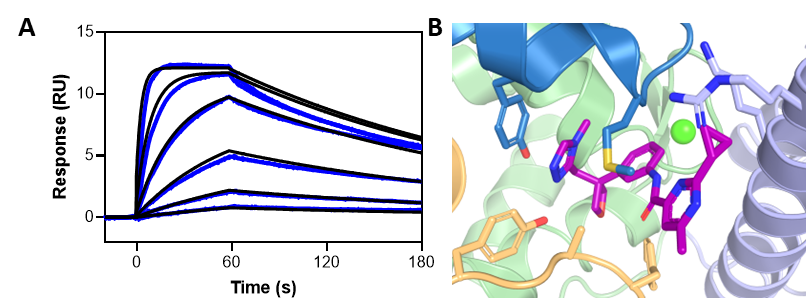
